# Integration of metagenome-assembled genomes with clinical isolates reveals genomic signatures of *Klebsiella pneumoniae* in carriage and disease

**DOI:** 10.1101/2024.12.17.628241

**Authors:** Samriddhi Gupta, Alexandre Almeida

**Affiliations:** Department of Veterinary Medicine, University of Cambridge, UK

## Abstract

*Klebsiella pneumoniae* is an opportunistic pathogen causing diseases ranging from gastrointestinal disorders to severe liver abscesses. While clinical isolates of *K. pneumoniae* have been extensively studied, less is known about asymptomatic variants colonizing the human gut across diverse populations. Genome-resolved metagenomics has offered unprecedented access to metagenome-assembled genomes (MAGs) from diverse host states and geographical locations, opening opportunities to explore health-associated microbial features. Here we analysed 662 human gut-derived *K. pneumoniae* genomes (319 MAGs, 343 isolates) from 29 countries to investigate the population structure and genomic diversity of *K. pneumoniae* in carriage and disease. Only 9% of sequence types were found to be shared between healthy and disease states, highlighting distinct diversity across health conditions. Integrating MAGs nearly doubled gut-associated *K. pneumoniae* phylogenetic diversity, and uncovered 86 lineages without representation among >20,000 *Klebsiella* isolate genomes from various sources. Genomic signatures linked to pathogenicity and carriage included those involved in antibiotic resistance, iron regulation, restriction modification systems and polysaccharide biosynthesis. Notably, machine learning models integrating MAGs and isolates more accurately classified disease and carriage states compared to isolates alone. These findings showcase the value of metagenomics to understand pathogen evolution with implications for public health surveillance strategies.

## Main

The species *Klebsiella pneumoniae* is a gram-negative, facultative anaerobic opportunistic pathogen found in the human upper respiratory and intestinal tracts^1^. Historically, *K. pneumoniae* strains were classified based on their capsular types (K-types) and lipopolysaccharide O-antigen structures. However, the use of genetic markers and the wider adoption of whole genome sequencing led to an improvement in the characterization of *K. pneumoniae* diversity. Diancourt et al.^2^ introduced a multi-locus sequencing typing (MLST) scheme based on seven housekeeping genes, enabling the categorisation of *K. pneumoniae* populations into distinct sequence types (STs). These STs have been instrumental in understanding the genotypic and phenotypic diversity of *K. pneumoniae* subspecies, especially in the context of their virulence potential. Based on the presence of specific virulence genes, *K. pneumoniae* can be divided into classical (cKP) and hypervirulent (hvKP) strains. A previous study^3^ identified five genotypic markers that effectively differentiate hvKP from cKP strains: *iucA* (aerobactin siderophore biosynthesis), *iroB* (salmochelin siderophore biosynthesis), *peg-344* (putative transporter) and *rmpA/rmpA2* (regulators of capsule production). cKP strains are also more commonly associated with two major antimicrobial resistance (AMR) mechanisms: the production of extended beta-lactamases (ESBLs) and carbapenemases. These resistant strains have globally disseminated and, in 2019, *K. pneumoniae* was identified as one of the six pathogens responsible for over 250,000 deaths associated with AMR^4^. Consequently, the World Health Organization has classified carbapenem-resistant *Enterobacteriaceae*, including *K. pneumoniae*, as priority pathogens in urgent need of new treatments.

While significant advances have been made through genomic studies of *K. pneumoniae* clinical isolates, studies on human carriage strains circulating asymptomatically in the global population are limited. Although *K. pneumoniae* is part of the human gut microbiome, carriage strains may acquire pathogenic traits that further increase the risk of infection^5,6^, especially among immunocompromised individuals^7^. *K. pneumoniae* strains colonising the gut microbiome have also been linked to gastrointestinal disorders such as inflammatory bowel disease^8^, colorectal cancer^9^ and diarrhoea^10^. Furthermore, it has been suggested that both symptomatic and asymptomatic colonization of the intestinal tract serves as a major reservoir for transmission to other sterile sites^11^ and cause extraintestinal infections such as urinary tract infections (UTIs), bacteraemia, liver abscesses, meningitis, endophthalmitis and osteomyelitis. For instance, a prior study on a cohort of 498 patients found that *K. pneumoniae* gut colonization on admission was significantly associated with subsequent extraintestinal infections such as pneumonia, wound infections, UTIs and bacteraemia with sepsis^12^. Altogether, these studies demonstrate that gut colonization is a major risk factor for nosocomial *K. pneumoniae* infections, highlighting the need for research efforts investigating the pathogenic potential of gut-associated lineages.

Developments in metagenomic methods have transformed our understanding of the diversity of the species, strains and genes found within the human gut microbiome^13–16^. In metagenomics, the entirety of the genomic content of a sample is purified, fragmented and sequenced. The resulting DNA segments can then be assembled into contigs to reconstruct the genomes of the microorganisms present in the sample, which are referred to as metagenome-assembled genomes (MAGs)^17^. Large sequence catalogues derived from the human gut microbiome are now available in public databases such as the Unified Human Gastrointestinal Genome (UHGG)^18^ catalogue. These include genomes recovered from faecal samples of various host states and geographic locations, providing access to diverse collections of microbial lineages naturally found in the human population.

In this study, we compiled 662 high-quality MAGs and isolates of gut-derived *K. pneumoniae* from 29 countries to explore their diversity and genomic properties across distinct health states. Our results highlight the value of integrating MAGs in pathogen genomic studies to identify diagnostic markers of carriage and disease, and infer the relative infection risk of human gut-colonized lineages.

## Results

### Genotyping *K. pneumoniae* in carriage and disease

To investigate the genetic features of carriage and clinical *K. pneumoniae* isolates from the human gut, we first compiled all high-quality MAGs (*n* = 319) and isolate genomes (*n* = 343) from the Unified Human Gastrointestinal Genome (UHGG) catalogue (Supplementary Table 1). The UHGG represents a comprehensive collection of isolates and MAGs from the human gut microbiome derived from >11,000 metagenomic samples worldwide. We collected and curated additional genome metadata regarding the health status and country of origin from their associated samples. Of the 662 genomes, health status information was obtained for 527 genomes, with 132 being classified as carriage- and 395 as disease-associated genomes. With regards to geographical distribution, the *K. pneumoniae* genomes spanned 29 countries, with the majority originating from China (*n* = 207) and the United States (*n* = 167). Overall, 511 genomes had complete metadata for both health status and country of origin (Fig. 1a).

**Figure 1.**
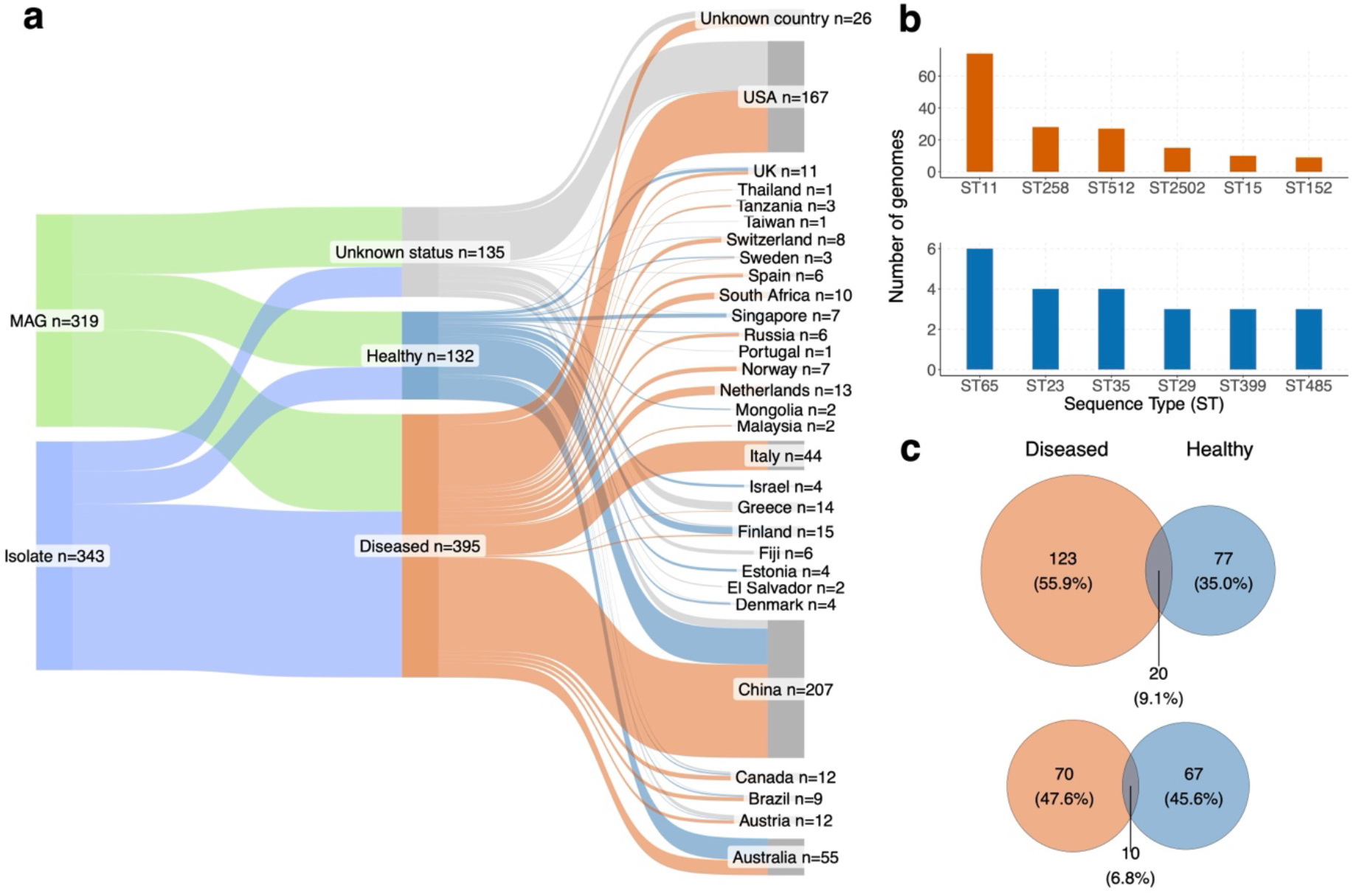
Global collection of *K. pneumoniae* MAGs and isolates. **a**, Metadata distribution of the 662 *K. pneumoniae* genomes analysed. Data is partitioned into three metadata factors: genome type (metagenome-assembled genomes, MAGs or isolates), health status (diseased, healthy or unknown) and country of origin. **b**, Sequence Types (STs) of the faecal-derived *K. pneumoniae* among disease (top) and carriage (bottom) strains. Only the top six STs among each group are shown. **c**, Venn diagram showing the intersection between STs detected among diseased and carriage genomes across all countries (top) or only considering countries where both carriage and disease genomes were recovered (bottom).

We assessed the distribution of sequence types (STs) derived from Kleborate^19^ according to the health status of the colonized host (Fig. 2b). A total of 220 STs were assigned to the 527 genomes with health status information. Among the disease-associated genomes, three *K. pneumoniae* carbapenemase (KPC)-producing lineages were the most prevalent: ST11 (*n* = 74), followed by ST258 (*n* = 28) and ST512 (*n* = 27). Other STs contained fewer than 16 disease-associated genomes. In contrast, carriage strains were more evenly distributed, with ST65, ST23, ST35 and ST29 being the most frequent, but only represented by a maximum of 6 genomes (Fig. 1b). Comparing the lineages shared across health states showed that only 9% of STs (20/220) were assigned to both carriage and disease genomes (Fig. 2c). In contrast, 123 STs (56%) were unique to disease lineages, whereas 77 (35%) were exclusive to carriage. To account for geographical biases, we further compared the level of ST overlap by only considering countries with genomes obtained from both carriage and disease (Fig. 2c), which further confirmed limited sharing between health states (7%). Overall, this shows that carriage and disease populations of *K. pneumoniae* in the human gut are mostly genotypically distinct.

**Figure 2.**
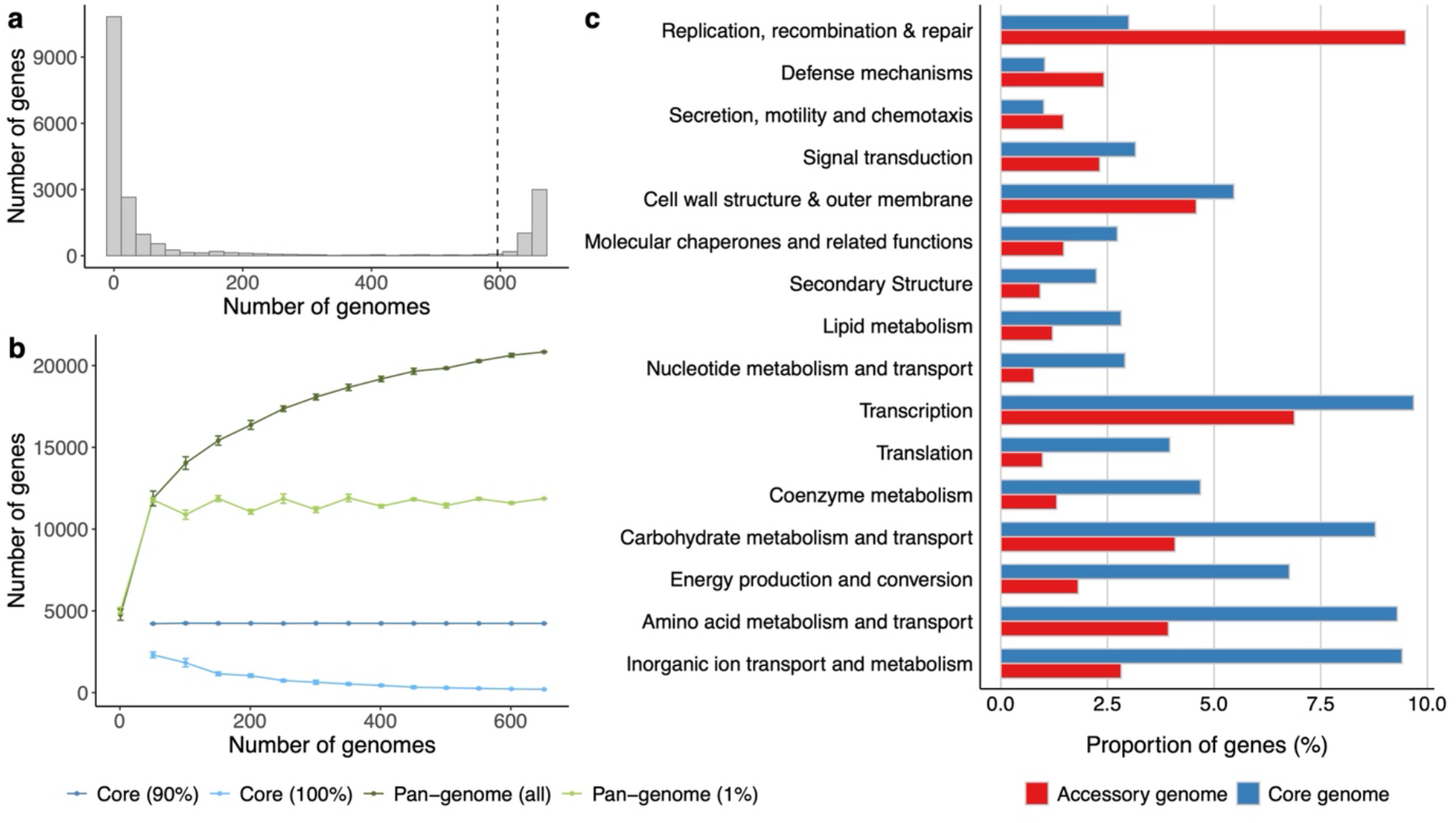
Pan-genome patterns of *K. pneumoniae*. **a,** Distribution of the number of genes detected according to the number of *K. pneumoniae* genomes they were found in. A threshold of 90% (vertical line) was used to define core genes. **b,** Core- and pan-genome accumulation curves obtained according to different filtering thresholds. **c,** Functional categories differentially abundant between the accessory and core genome, tested with a two-sided Fisher’s exact test (adjusted *P* < 0.05).

### Pan-genome patterns and functions of *K. pneumoniae*

To explore the core and accessory genome of the collection of gut-derived *K. pneumoniae*, we used the pan-genome analysis tool Panaroo^20^ tested with various configurations (Extended Data Fig. 1). Across different parameters, we obtained a mean pan-genome size of 21,160 genes (interquartile range, IQR = 20,738–21,559) and 4,117 core genes (IQR = 4,050–4,182). Core- and pan-genome estimates were generally consistent across different settings, with maximum variations in pan-genome size and number of core genes of 5% and 3%, respectively. Downstream analysis were performed using the moderate filtering mode, 90% identity and non-merged paralogs to increase overall recall and mitigate the risk of technical artefacts leading to multiple variants of the same gene.

Using the above criteria, we then characterized pan-genome patterns of *K. pneumoniae* based on gene frequency and pan-genome accumulation curves (Fig. 2). We observed a clear separation of core and accessory genes using a 90% threshold of gene presence (Fig. 2a). Although the pan-genome curve using all accessory genes suggested an open pan-genome for *K. pneumoniae*, this was not the case when only considering genes found in at least >1% of the genomes (Fig. 2b). This suggests that newly acquired genes are very rare within the whole *K. pneumoniae* population. Similarly, a 90% threshold for defining core genes revealed a more consistent trend compared to using a more strict 100% cut-off (Fig. 2b).

Annotation of the pan-genome highlighted functional differences between the core and accessory genomes (Fig. 2c). Accessory genes were significantly overrepresented (two-sided Fisher’s exact test, *P* < 0.05) in functions related with replication, recombination and repair, as well as defence mechanisms. In contrast, core genes were predominantly associated with inorganic ion and amino acid metabolism, in addition to energy production. Of note, 61% of the accessory and 23% of the core genome of *K. pneumoniae* could not be assigned to a known functional category, highlighting the extent of uncharacterized genetic diversity still to be explored even in well-known species of clinical relevance.

### Metagenome-assembled genomes expand *K. pneumoniae* diversity

Using the core genes identified with Panaroo, we investigated the phylogenetic relationship of the *K. pneumoniae* MAGs and isolate genomes here analysed. The phylogenetic tree built from the core gene alignment exhibited a well-defined structure consistent with the predicted ST, with MAGs and isolates largely interspersed across the phylogeny (Fig. 3a). Considering the mixture of MAGs and isolate genomes could lead to increased technical variation, we compared the inferred population structure using different analysis approaches (Extended Data Fig. 2). We compared three clustering methods: i) the gene-based core tree obtained with Panaroo; ii) a SNP-based core tree generated by Snippy^21^ with recombination removed; and iii) gene presence/absence patterns derived from the pan-genome data. There was a strong correlation between the three approaches tested (Mantel test, Pearson’s r = 0.77–0.91; *P* < 0.0001), showing that the clustering structure was consistent across core genes, SNPs and the whole pan-genome (Extended Data Fig. 2). We then assessed the relationship between health state, geographical location, genome type (MAG or isolate) and source (community or hospital) with the aforementioned three clustering measures (Fig. 3b). Statistical analysis using a PERMANOVA test revealed that the specific health state (disease name or condition) and country of origin had the strongest association (*P* < 0.001) with the population structure of *K. pneumoniae* (Fig. 3b).

**Figure 3.**
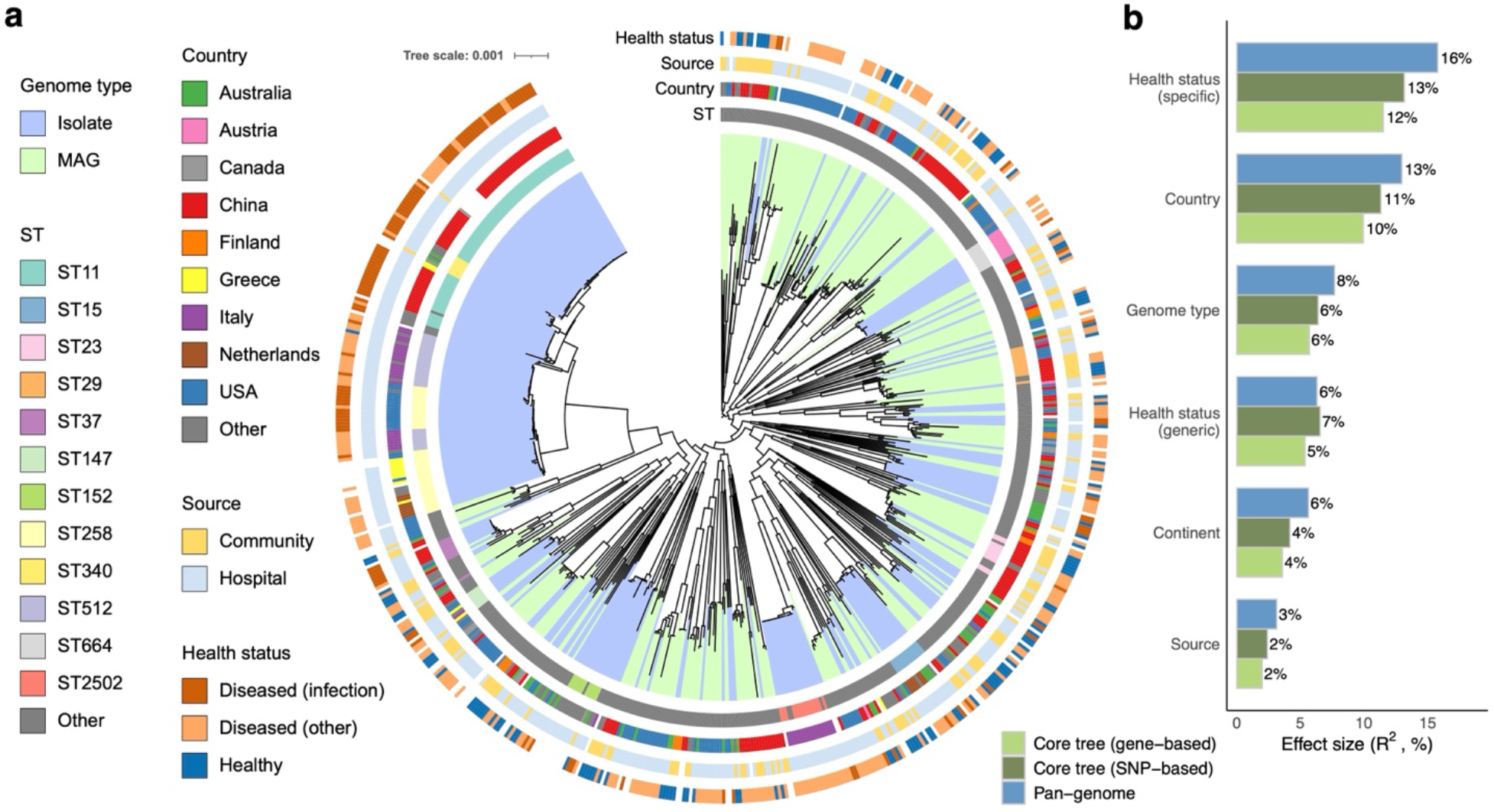
Phylogenetic diversity of *K. pneumoniae* MAGs and isolates. **a,** Phylogenetic relationship of the dataset of 662 faecal-derived *K. pneumoniae* genomes based on phylogenetic distances of the gene-based core tree obtained with Panaroo. **b**, Effect of various metadata variables on the population structure of faecal-derived *K. pneumoniae*, measured by the pairwise cophenetic distances using either the core trees (gene- or SNP-based) or the pan- genome distances (Jaccard). A PERMANOVA test was used to quantify the effect size (R^2^) and statistical significance. All associations were found to be statistically significant (*P* < 0.001).

As MAGs can uncover new lineages not represented by clinical isolates, we further quantified how much genetic diversity within the gut-derived *K. pneumoniae* population was represented by each genome type. We performed a phylogenetic diversity (PD) analysis using Faith’s PD method, which showed that the inclusion of MAGs provided a 88% and 96% increase in phylogenetic diversity among disease and carriage lineages, respectively. Of note, we did not identify any MAGs belonging to the disease-associated clade dominated by ST11/ST258 isolates (Fig. 3). Given that >11,000 faecal metagenomic samples are represented within the UHGG, their absence from the catalogue suggests these lineages are very rare among asymptomatic carriers of *K. pneumoniae*. To further understand whether the MAGs here included matched *K. pneumoniae* isolate genomes from other sources, we conducted a comparative genomic analysis using Mash against all isolate genomes from the *Klebsiella* genus available on the NCBI RefSeq database (*n* = 20,792 genomes). Comparison of the 319 gut-derived *K. pneumoniae* MAGs against the RefSeq genomes showed a median distance of 0.0034 (interquartile range, IQR = 0.0024–0.0052) with their closest matches (Supplementary Table 1 and Extended Data Fig. 3a). Interestingly, the Mash distances exhibited a bimodal distribution separated between a distance threshold of 0.005 (approximately equivalent to 99.5% average nucleotide identity), potentially reflecting a strain-level boundary. We detected 86 MAGs in particular with Mash distances greater than 0.005 to their closest RefSeq sequence, which therefore suggests these represent understudied lineages without a reference isolate genome. These MAGs spanned distinct health states and geographical regions (Extended Data Fig. 3b), illustrating the value of using genome-resolved metagenomics for uncovering a more complete view of pathogen diversity and evolution.

### Genomic signatures of carriage and disease

Having a collection of MAGs and isolates across diverse health states provided the opportunity to investigate genetic features associated with health and disease. First, we evaluated virulence and antibiotic resistance profiles of *K. pneumoniae* estimated with the Kleborate^19^ tool (Fig. 4 and Supplementary Table 2). In particular, we performed a statistical analysis to determine whether virulence and resistance scores differed between disease- and carriage-associated genomes (Fig. 4a). Although virulence scores were not significantly different (Wilcoxon rank-sum test, *P* = 0.516), resistance scores were significantly higher in disease-associated strains (*P* < 0.0001). The most prevalent resistance determinants were the mutations 35Q responsible for resistance to penicillin and cephalosporins^19^; ParC-80I and GyrA-83I involved in fluoroquinolone resistance^22,23^; a β-lactamase gene coding for TEM-1D.v1^19,24^; the *sul1* gene associated with resistance to sulfonamides^25^; and the carbapenemase variant KPC-2^26^ (Fig. 4b). Therefore, these results suggest that antibiotic resistance profiles, as opposed to virulence genes, provide a better distinguishing signature of different host states.

**Figure 4.**
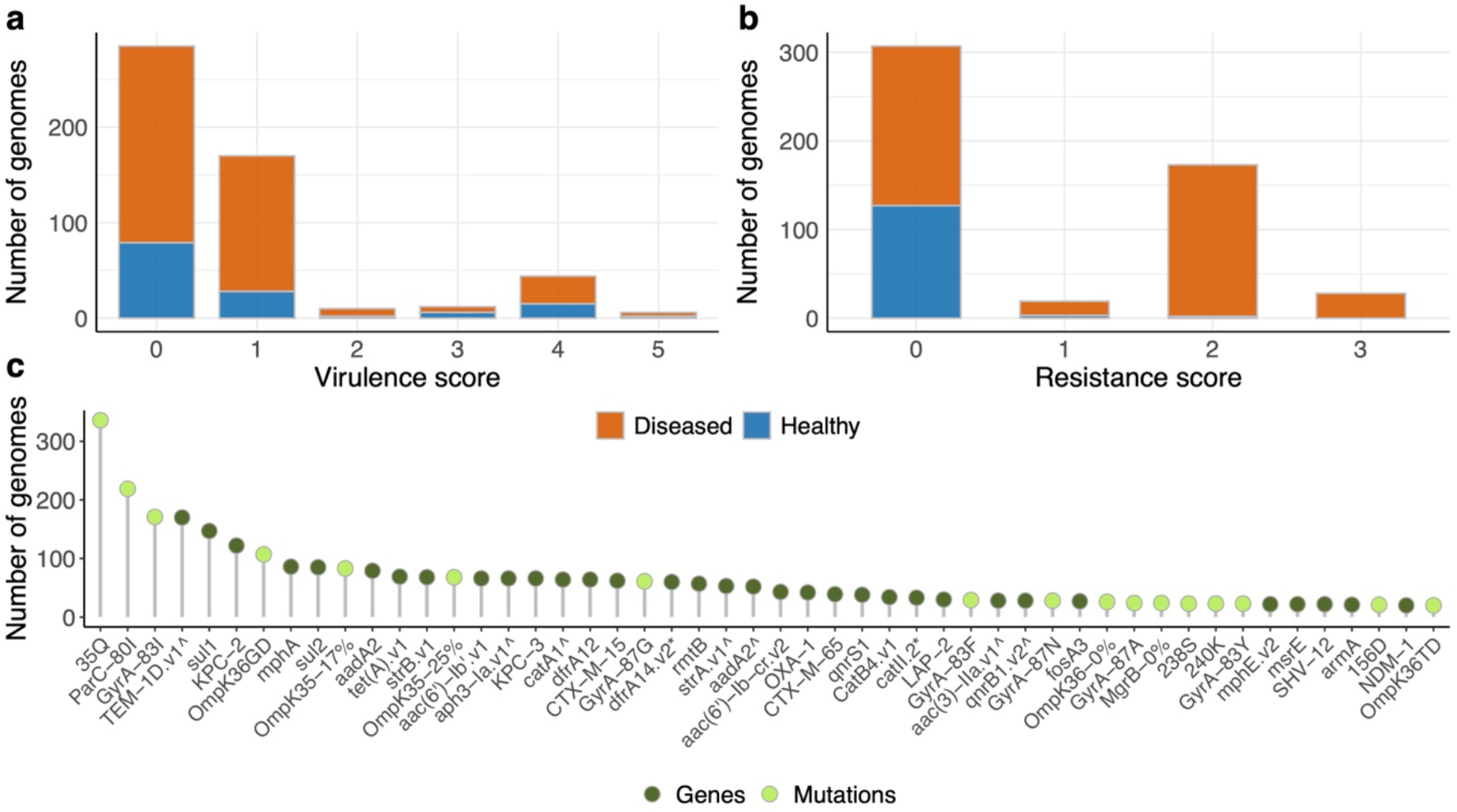
Virulence and resistance patterns of carriage and disease lineages. **a**, Distribution of virulence scores according to the health state of the sampled individual. The virulence scores are given as follows: 0 = None; 1 = *ybt*; 2 = *ybt* + *clb*; 3 = *iuc*; 4 = *ybt* + *iuc*; 5 = *ybt* + *clb* + *iuc*. **b**, Distribution of resistance scores based on the following criteria: 0 = ESBL-, Carb-; 1 = ESBL+, Carb-; 2 = Carb+; 3 = Carb+, Col+**. c**, Top 50 antimicrobial resistance determinants (genes or mutations) detected across the whole *K. pneumoniae* genome collection analysed.

Next, to extend our analyses more broadly to all genetic features associated with health status among *K. pneumoniae*, we performed a microbial Genome Wide-Association Study (mGWAS) at both gene and SNP level. To account for inherent genomic differences between MAGs and isolate genomes, we included ‘genome type’ as a covariate in the models, and further excluded any genes found to be significantly associated with either MAGs or isolates (see Methods for further details). In the end, we identified 484 genes significantly different according to health status (FDR < 5%, Supplementary Table 3): 344 associated with infection or carriage (Fig. 5 and Extended Data Fig. 4), and 181 from the analysis of all disease lineages. At the SNP level, only one mutation (CP012745:g.1749711C>T) was found to be significantly overrepresented among genomes from infection. This SNP was located in an intergenic region between two glutathione S-transferase genes. Glutathione S-transferases are a family of enzymes involved in the detoxification of endogenous and xenobiotic compounds^27^.

**Figure 5.**
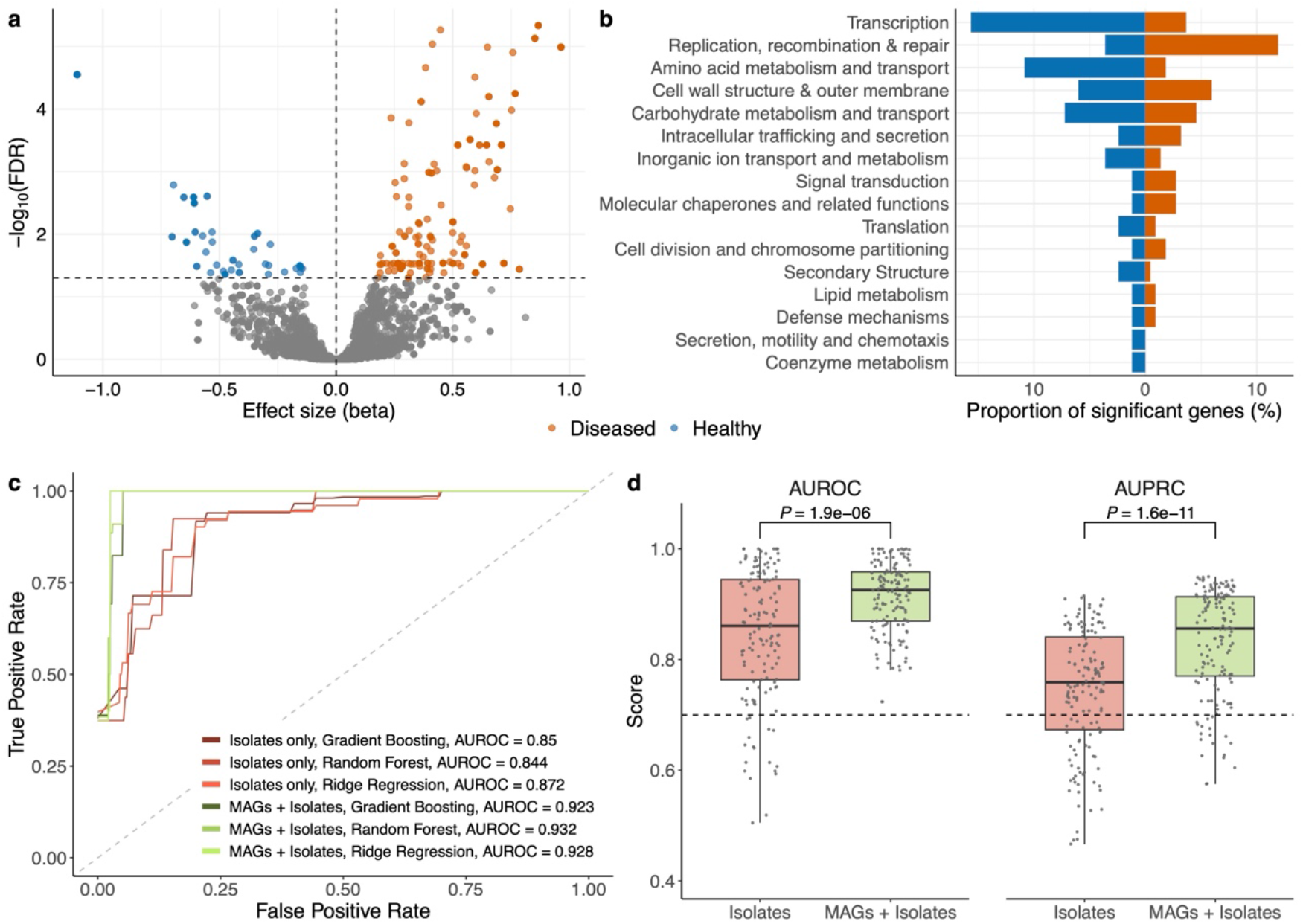
Genomic signatures of *K. pneumoniae* in carriage and infection. **a**, Distribution of effect sizes (beta) and -log10–transformed adjusted *P* values (False Discovery Rate, FDR) of genes significantly present or absent among infection-associated *K. pneumoniae* genomes. **b**, Proportion of the significant genes assigned to different COG categories. Orange bars to the right indicate the number of genes overrepresented in disease (infection) lineages, while the blue bars show genes overrepresented in carriage. **c**, Receiver operating characteristic (ROC) curve of the machine learning results linking the pan-genome patterns with the health state of the individual sampled. **d**, Comparison of Area Under the Receiver Operating Curve (AUROC) and Area Under the Precision-Recall Curve (AUPRC) scores of machine learning (ML) models classifying carriage and disease lineages using isolates alone or MAGs together with isolate genomes. Each box represents the interquartile range (IQR) across 150 independent seeds (50 per model). The centre line within the box represents the median score. Whiskers are shown extending to the furthest point within 1.5 times the IQR from the box. *P* values were derived from a two-sided Wilcoxon rank-sum test.

At the gene level, we focused on the candidate genes specifically associated with infection or carriage, as these are potentially more directly linked to the pathogenesis of the disease. Grouping the 344 significant genes into Clusters of Orthologous Groups (COGs) showed that categories involved in transcription and amino acid metabolism were overrepresented in carriage (two-sided Fisher’s exact test, FDR < 5%), while DNA replication and repair were more predominant in disease (at an FDR = 12%, Fig. 5b). However, 210 genes could not be assigned to a known function, representing a large genetic repertoire that may underly uncharacterized disease mechanisms. Among the most significant carriage genes with a predicted function was a genetic cluster containing a transcriptional repressor similar to the Ferric uptake regulator (Fur) family. Members of the Fur family have been shown to repress siderophore activity which in turn can reduce *K. pneumoniae* virulence^28,29^. An overrepresentation of these genes among carriage strains suggests they may attenuate the pathogenic potential of *K. pneumoniae*. In contrast, infection-associated genes with the highest effect size were related with restriction modification (RM) systems, phage proteins and polysaccharide biosynthesis. Although surface polysaccharides have been previously implicated in *K. pneumoniae* infection^29^, the role of RM genes and phages in disease has been less studied. RM systems are responsible for bacterial protection against foreign DNA, such as that encoded by bacteriophages^30^. We hypothesize that the ability of these lineages to resist infection by certain bacteriophages may provide a fitness advantage compared to carriage strains.

To further assess how well the pan-genome diversity was able to distinguish carriage from infection-associated lineages, we trained supervised machine learning models (ridge regression, random forest and gradient boosting) to classify *K. pneumoniae* genomes based on the health status (infected or healthy) of their hosts. All models showed good classification performance (Area Under the Receiver Operating Curve, AUROC >0.9; Fig. 5c and Extended Data Fig. 5), even by broadening the analysis to all disease lineages (median AUROC >0.8; Extended Data Fig. 5). Notably, models trained with both MAGs and isolates performed significantly better (two-sided Wilcoxon rank-sum test, *P* < 0.001) than those with isolates alone, especially when evaluating model performance with the Area Under the Precision-Recall Curve (AUPRC, Fig. 5d). Therefore, these results show that combining both MAGs and isolates significantly improves the ability to distinguish genomic signatures related with health and disease.

## Discussion

Pathogen genomic studies have predominantly focused on the analysis of clinical isolates to derive biological insights into bacterial pathogenesis. In this study, we integrate over 600 metagenome-assembled genomes (MAGs) with clinical isolate genomes of *K. pneumoniae* from both carriage and disease states to improve our understanding of their evolution, phylogeography, and health-associated features. We focused primarily on genomes from the human gut given the importance of gut colonization as a risk factor for *K. pneumoniae* infections^1,11,12^. Importantly, we highlight that the inclusion of MAGs not only expanded the diversity of *K. pneumoniae*, but also substantially improved the ability to distinguish carriage from clinical lineages. Furthermore, we found that out of the available metadata linked to each genome, the specific health state and country of origin had the strongest association with the population structure of *K. pneumoniae*. Our findings expand on previous work investigating the global genomic diversity of *K. pneumoniae* in humans and animals^29^ across different health states.

Comparative analysis of carriage and disease genomes showed that disease-associated lineages were dominated by three KPC-producing strains — ST11, ST258 and ST512. ST11 has been recognized as one of the leading clinical pathogens to cause outbreaks in China^31^ while ST258 is commonly found in the United States and several other countries, including Canada and Italy^32^. ST512 has been located in various countries such as Finland, Italy and recently Algeria^33–35^. In contrast, asymptomatic carriage strains showed a more diverse distribution across sequence types, which were largely distinct from those assigned to disease lineages. By profiling the genetic markers involved in antibiotic resistance and virulence, we observed that higher resistance scores were a prominent characteristic of disease-associated genomes. This aligns with a previous genomic study in Northern Italy which showed that species and strains outside the hospital environment exhibited low levels of resistance^36^.

Our mGWAS analysis revealed 484 genes that were significantly different between health states, including those involved in transcriptional regulation and restriction modification systems that have not been previously implicated in *K. pneumoniae* pathogenesis. Moreover, machine learning models trained on the pan-genome were shown to accurately differentiate infection and carriage strains. Overall, these results show that disease-associated lineages harbour unique genetic traits that may confer increased pathogenesis to *K. pneumoniae* and serve as potential diagnostic markers.

Although we reveal important insights into the genomic features of carriage versus disease-associated *K. pneumoniae*, it is worth noting some of our study’s limitations. Despite our strict filtering criteria, MAGs should still be treated with caution compared to isolate genomes due to a higher risk of contamination from closely-related species^37^. Furthermore, even though we analysed a globally diverse genome collection, our samples were overrepresented in certain well-studied regions such as China and the United States. Lastly, given the genomic focus of our study, the uncharacterized candidate genes here identified will inherently require further experimental testing to explore their mechanistic role in the aetiology of disease.

In summary, our work underscores the importance of integrating MAGs with clinical isolates for population-based genomic analyses, identifying specific features of *K. pneumoniae* that could serve as markers of carriage and disease. Ultimately, this could lead to the development of targeted public health measures and therapeutic approaches aimed at reducing infection risk.

## Acknowledgements

The authors thank Ana Catarina da Silva for assisting in the curation of the sample metadata. We also thank Sebastian Bruchmann, Efrat Muller and all members of the Microbiome Function and Diversity group for helpful feedback and suggestions. Funding was provided by a Career Development Award from the Medical Research Council (MR/W016184/1) to A.A.

## Author contributions

S.G. performed the genomic analyses and wrote the manuscript draft. A.A. supervised the work, assisted in the analyses and edited the manuscript.

## Methods

### Genome collection and quality control

We first extracted all *Klebsiella pneumoniae* genomes derived from faecal samples available in the Unified Human Gastrointestinal Genome (UHGG)^18^ v1.0 database (*n* = 985). This dataset comprised two genome types: metagenome-assembled genomes (MAGs) and isolates. Given MAGs are generally of lower quality than isolate genomes, we performed strict quality control procedures. First, we applied a filter of >90% completeness and <5% contamination based on the genome statistics obtained with CheckM^38^ v1.0.11. Thereafter, we used GUNC^39^ v1.0.3 to further exclude genomes with both a ‘clade_separation_score’ >0.45 and ‘contamination_portion’ >0.05. Lastly, all genomes underwent taxonomic verification for *K . pneumoniae* using Kleborate^19^ v2.4.1 using default parameters. The final filtered dataset comprised a total of 662 genomes (319 MAGs and 343 isolate genomes; Supplementary Table 1).

### Metadata curation and genotyping

Metadata regarding health status, country of origin and source for each genome was gathered from the National Center for Biotechnology Information (NCBI) or the European Nucleotide Archive (ENA). Furthermore, a review of associated research papers (if available) was conducted to supplement the metadata obtained. Individuals classified as diseased were divided into two groups: i) those with conditions directly associated with or caused by the colonization of pathogenic *K. pneumoniae* strains, including infections and diarrhoeal diseases; or ii) any other diseases such as colorectal cancer, autoimmune disorders, liver disease, among others. Carriage strains were defined as those obtained from individuals explicitly classified as healthy. Sequence Types (STs) for each genome were derived with Kleborate, which performs sequence alignment against an established MLST scheme^2^ comprising seven loci (more information can be found in the associated BIGSdb resource: https://bigsdb.pasteur.fr/klebsiella/).

### Pan-genome analysis

We constructed a pan-genome of the 662 *K. pneumoniae* genome*s* selected using Panaroo^20^ v1.3.3, which enabled us to differentiate between the core and the accessory genes. We compared the use of different parameters to assess the impact of varying stringency levels in identifying the core genome. Specifically, Panaroo was run with ‘mode’ moderate and strict, which required each gene to be present in at least 1% or 5% of the genomes, respectively. For each mode configuration, we also tested a sequence identity threshold of both 90% and 95%. Additionally, the approach of handling paralogous genes was compared between merging and keeping them separate. The core genome threshold was set to 90%.

Functional analysis of the core and accessory genomes was performed using eggNOG- mapper^40^ v2.1.3. Genes were classified based on their assigned Cluster of Orthologous Group (COG) functional category. The percentage of genes in each COG category was calculated separately for both core and accessory genomes.

### Phylogenetic analyses

We used Snippy^21^ v4.6.0 (18) to extract single nucleotide polymorphisms (SNPs) of each *K. pneumoniae* genome against reference GUT_GENOME147590 (Supplementary Table 1), representing a finished isolate assembly. Gubbins^41^ v3.3.1 was utilized to identify and remove regions of recombination from the SNP alignment file and to construct a filtered phylogenetic tree. In parallel, a gene-based phylogenetic tree was also built using the approximate maximum-likelihood method implemented in FastTree^42^ v2.1.11, based on the core gene alignment generated by Panaroo. Lastly, a pan-genome tree was derived by estimating pairwise Jaccard distances of gene presence/absence patterns across all genomes. The SNP-based and pan-genome trees were visually compared with the gene-based core tree from Panaroo through a tanglegram using the ‘tanglegram’ function of the ‘dendextend’ R package^43^. To construct the tanglegram, trees were converted to ultrametric distances (function ‘chronos’ ; ‘ape’ R package^44^) and rooted at their midpoints (function ‘midpoint.root’; ‘phylotools’ R package^45^). These were subsequently converted into dendograms (function ‘as.dendogram’; base R) and reordered to ensure that corresponding taxa were positioned similarly in both trees. A statistical analysis using the Mantel test was conducted (function ‘mantel’ ; ‘vegan’ R package^46^), using the Pearson correlation method with 999 permutations, to assess the correlation between the distances derived from each tree.

To investigate the association between the population structure of our dataset and key metadata factors (i.e., health status: diseased or healthy; genome type: MAGs or isolates; and country of origin of the samples), we performed a permutational multivariate analysis of variance (PERMANOVA) test using the ‘adonis2’ function from the ‘vegan’ R package^46^. As the ‘adonis2’ function requires a distance matrix as an input, the previously calculated cophenetic or Jaccard distances from the gene, SNP or pan-genome trees were used. The model formula specified the metadata factor of interest as the independent variable, with the distance matrix as the dependent variable.

To assess phylogenetic diversity (PD), we employed the Faith’s PD method, which quantifies diversity as the sum of all branch lengths. The phylogenetic tree constructed using Panaroo was rooted at midpoint and filtered to create a subset of the tree with isolates only. The increase in phylogenetic diversity provided by the MAGs was calculated as: (PD all – PD isolates) / PD isolates.

### Genomic comparison with public databases

To determine the similarity of gut-derived *Klebsiella pneumoniae* metagenome-assembled genomes (MAGs) to cultured *K. pneumoniae* genomes from other habitats, we performed a comparative genomic analysis using Mash^47^ v2.3 which calculates pairwise genome distances based on shared k-mer content. Each *K. pneumoniae* MAG was compared against a database of all cultured genomes from the *Klebsiella* genus available on NCBI RefSeq release 219 (*n* = 20,792 genomes). Thereafter, we identified the closest reference genome for each MAG based on the lowest Mash distance. A Mash distance threshold of 0.005 was used to determine which MAGs represented lineages without a reference isolate genome.

### Resistance and virulence gene analysis

Pathogenicity and antibiotic resistance profiles were calculated using Kleborate^19^ (Supplementary Table 2). This tool calculates a virulence and a resistance score for a given genome based on genotypic markers. The virulence scoring system ranges from 1 to 5, based on the following criteria: 1 = *ybt*, 2 = *ybt* + *clb*, 3 = *iuc*, 4 = *ybt* + *iuc*, and 5 = *ybt* + *clb* + *iuc*. The resistance score is calculated as follows: 1 = ESBL , 2 = carbapenemases, 3 = carbapenemases + colistin resistance gene. We assessed the association between virulence and resistance profiles and health status (diseased versus healthy) using a two-sided Wilcoxon rank-sum test.

### Microbial Genome Wide-Association Study (mGWAS)

Pyseer^48^ v1.3.11 was used to identify genetic features (genes and single nucleotide polymorphisms, SNPs) associated with health status. The gene presence/absence matrix generated with Panaroo was used as input for Pyseer to identify genes significantly present or absent in disease-associated *K. pneumoniae* genomes. However, to account for MAG incompleteness and/or contamination, only genes present in >1% and <90% of the population were analysed. Parallel to this, SNP data derived from Snippy were used to identify mutations associated with either carriage or diseased genomes.

mGWAS was performed solely on the genomes whose health status and country of origin was available. To account for population structure, the Fast-LMM (Factored Spectrally Transformed Linear Mixed Models) algorithm was utilized with a similarity kinship matrix. This was achieved using the script ‘phylogeny_distance.py’ from Pyseer with the ‘--lmm’ option to signify the use of a linear mixed model that considers the phylogenetic distance as a measure of genetic similarity. Country of origin and genome type (MAG or isolate) were additionally used as covariates in the mixed effect model. Statistical significance was determined after multiple testing correction using the Benjamini-Hochberg procedure and False Discovery Rate (FDR). An adjusted *P* value threshold < 0.05 was used to find genes and SNPs significantly associated with health or disease states (Supplementary Table 3). As an additional control, a separate mGWAS analysis was performed to identify genes and variants associated with genome type (MAG/isolate), and these were excluded from the disease analysis.

To elucidate the biological implications of our mGWAS findings, significant genes were annotated via eggNOG-mapper. The output from eggNOG-emapper included detailed descriptions of each gene and an associated functional category based on the Clusters of Orthologous Groups (COGs) classification. Uncharacterized genes were considered those either without an eggNOG annotation, or with an ‘S’ (Function Unknown) COG classification.

### Machine learning classification

We trained three supervised machine learning (ML) models (ridge regression, random forest and gradient boosting) to distinguish disease- from carriage-associated *K. pneumoniae* genomes. ML models were trained based on the pan-genome presence/absence data. Features were pre-processed to only include those found between 1% and 90% frequency. ML models were run using a custom workflow adapted from the ‘mikropml’ R package^49,50^ (https://github.com/alexmsalmeida/ml-microbiome). Only genomes whose metadata for both the health status and country of origin was available were utilised. Model training and hyperparameter tuning was performed on 80% of the data using a 5-fold cross-validation, while the other 20% were used for testing with the best hyperparameter setting. To evaluate model performance, each analysis was repeated with 50 unique random seeds. Additionally, the ST of the genomes was used as the grouping variable to account for population structure (i.e., genomes from the same ST were kept together in either the training or test set). Summary statistics (mean, median, interquartile range) for the Area Under the Receiver Operating Curve (AUROC) and the Area Under the Precision-Recall Curve (AUPRC) were used to evaluate model performance. Models trained with MAGs and isolates or isolates alone were compared to assess the diagnostic value of the MAGs for disease classification using a two-sided Wilcoxon rank-sum test. When performing model training with both MAGs and isolates, features found significantly associated with ‘genome type’ based on the mGWAS results were first excluded.

## Data availability

All genomes used are publicly available in the Unified Human Gastrointestinal Genome catalogue (https://ftp.ebi.ac.uk/pub/databases/metagenomics/mgnify_genomes/human-gut/v1.0/) and the European Nucleotide Archive (see Supplementary Table 1 for list of accession codes and genome identifiers). Pan-genome data files (gene presence/absence matrix and FASTA file) are available in: https://doi.org/10.6084/m9.figshare.27961089.v1.

## Code availability

Custom code and scripts are available in the following GitHub repository: https://github.com/microfundiv-lab/KpMAGs.

---

**Extended Data Figure 1.**
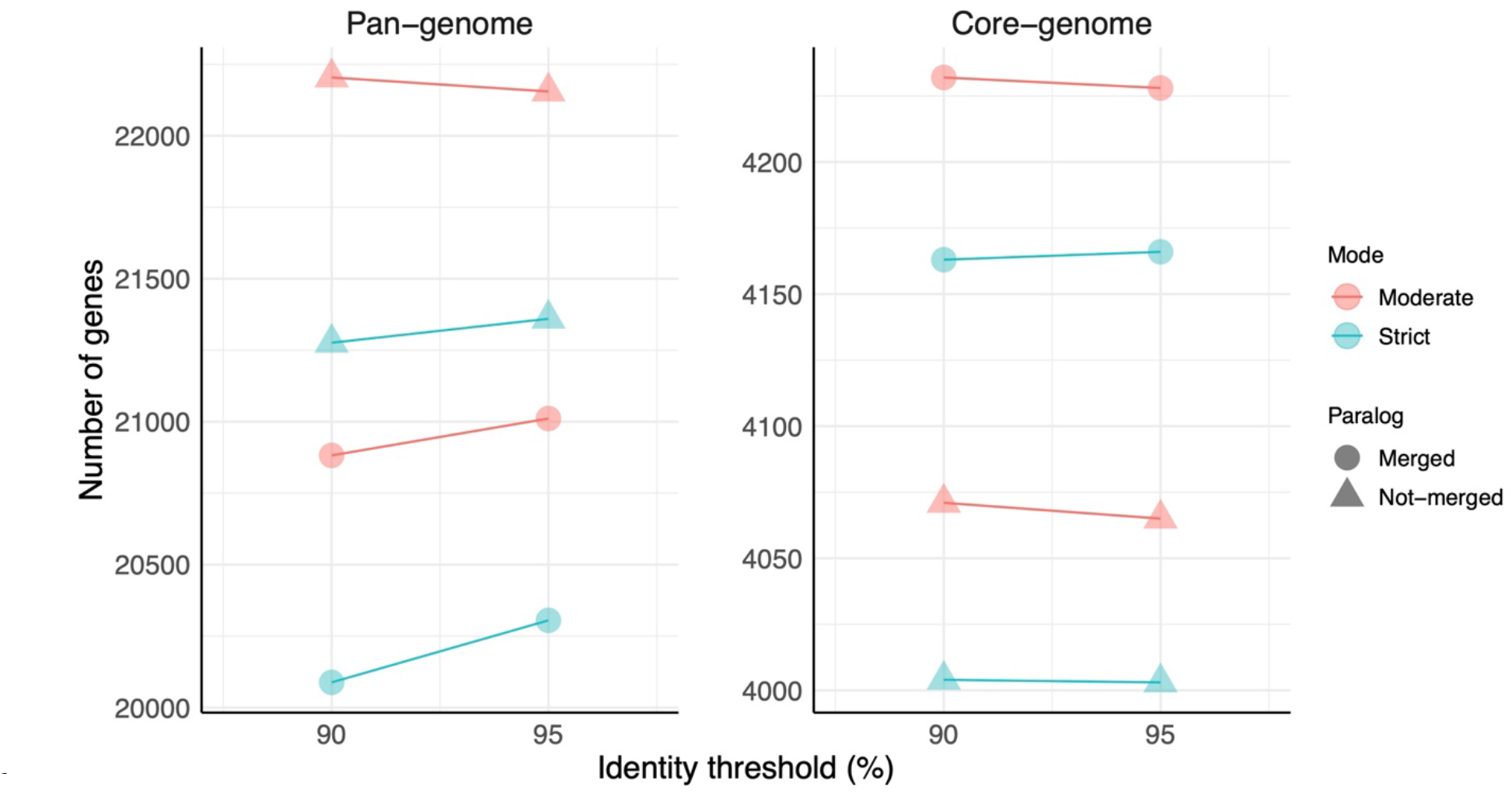
Pan-genome size and number of core genes identified with Panaroo. Variation in pan-genome size and total number of core genes identified in *K. pneumoniae* according to different parameters used within Panaroo. The Y-axis represents the number of genes and the X-axis the two sequence identity thresholds (90% or 95%). The colour denotes the mode used (i.e. strict or moderate) and the shapes the paralog handling method (merged or not merged). A core genome threshold was set at 90%.

**Extended Data Figure 2.**
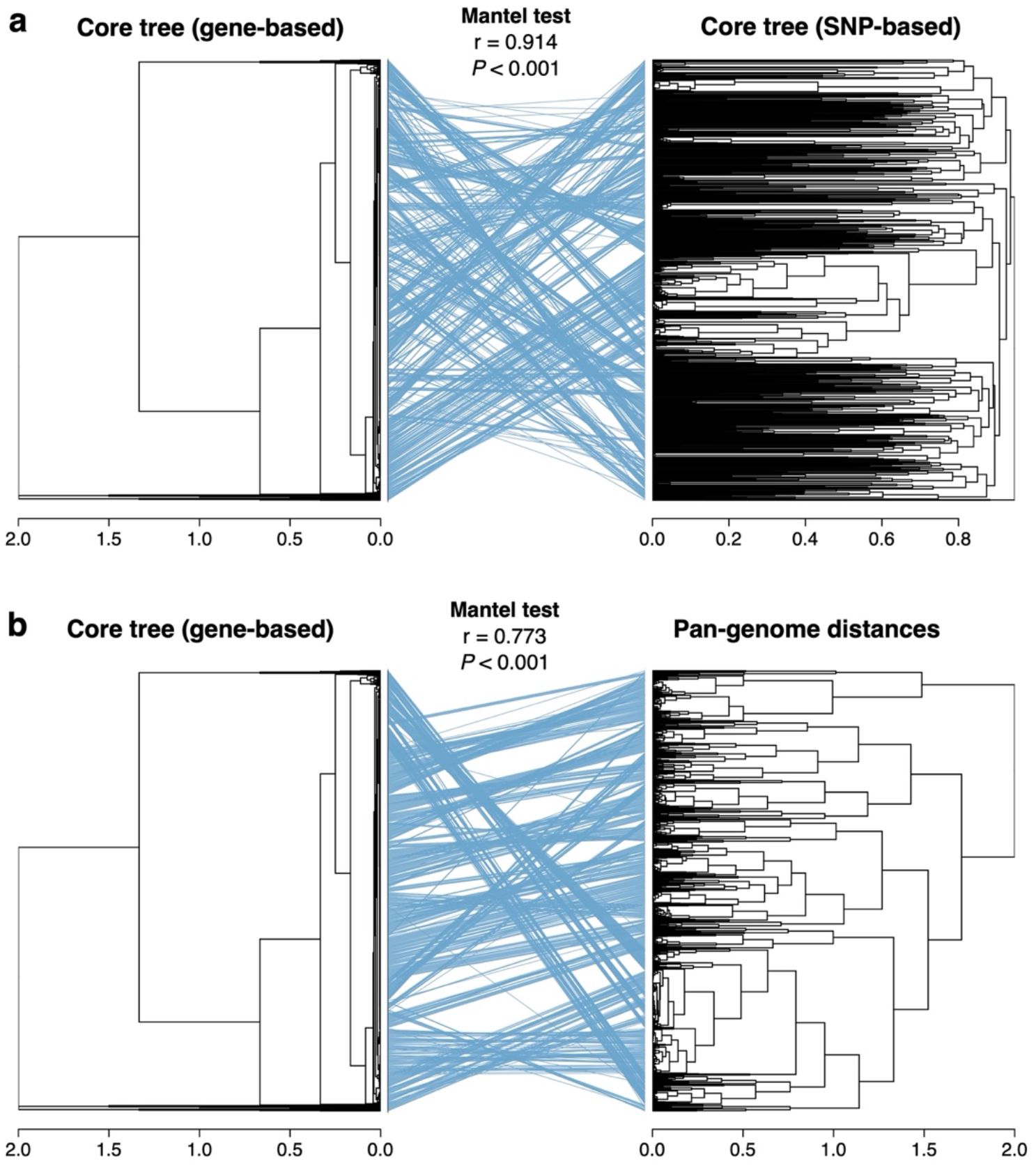
Correlation between phylogenetic clustering methods. **a,** Tanglegram comparing the gene-based core phylogenetic tree of *K. pneumoniae* generated by Panaroo against the SNP-based core tree obtained with Snippy. **b**, Comparison of the gene- based core phylogenetic tree against the genome clustering obtained with Jaccard pan- genome distances (gene presence/absence patterns). A Mantel test was used to assess the correlation between each pair of trees.

**Extended Data Figure 3.**
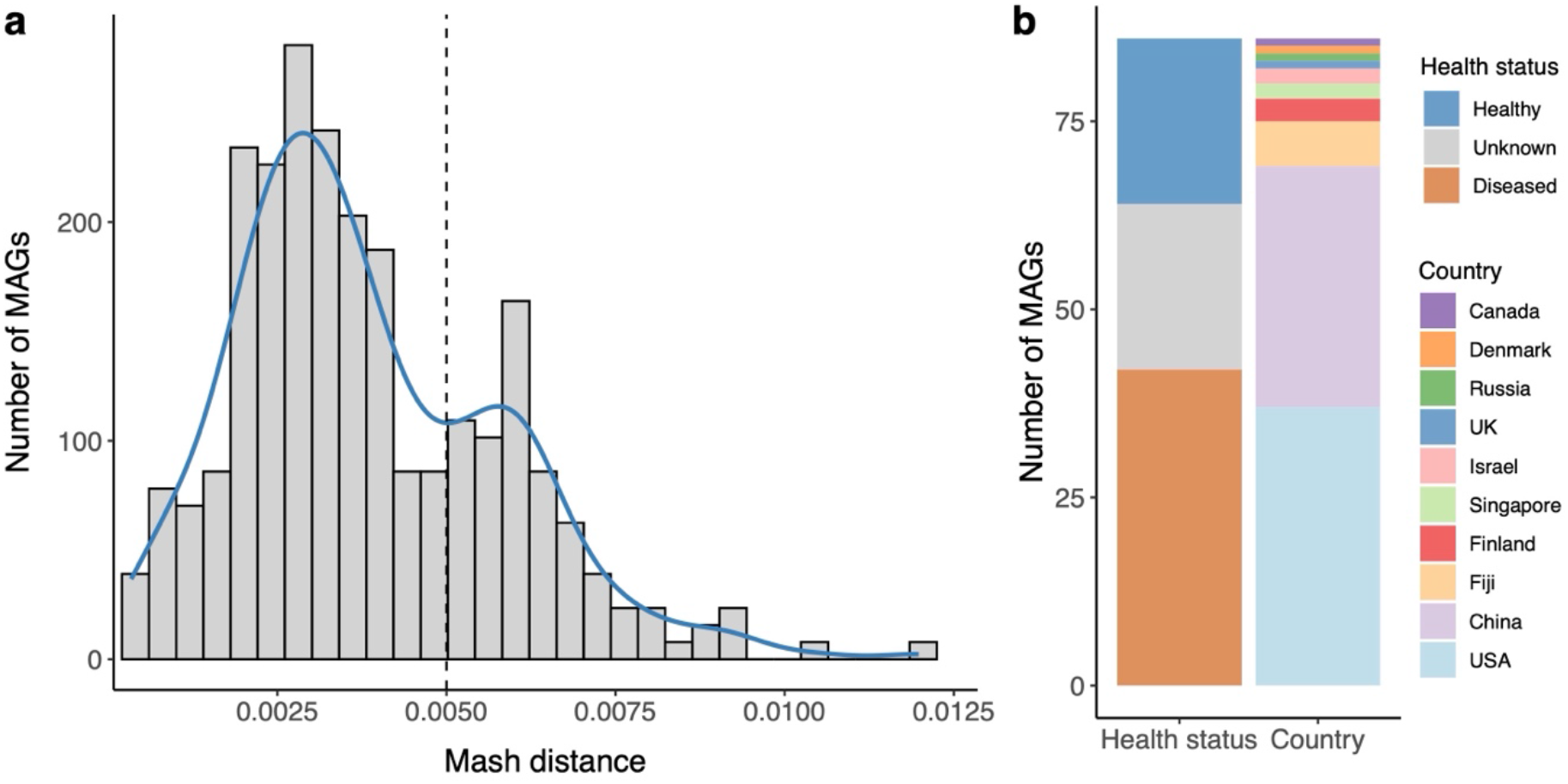
Comparison of metagenome-assembled genomes with reference isolate genomes. **a,** Distribution of Mash distances between the *K. pneumoniae* metagenome-assembled genomes (MAGs) and their closest reference genomes when compared against 20,792 *Klebsiella* isolate genomes from the NCBI RefSeq database. The histogram shows the frequency of MAGs at different Mash distance intervals, with a density curve (blue line) overlaid to illustrate the overall distribution. A threshold of 0.005 (vertical dashed line) was used to define lineages without a close reference genome. **b**, Metadata distribution of the 86 MAGs representing lineages without a reference isolate genome available on NCBI RefSeq.

**Extended Data Figure 4.**
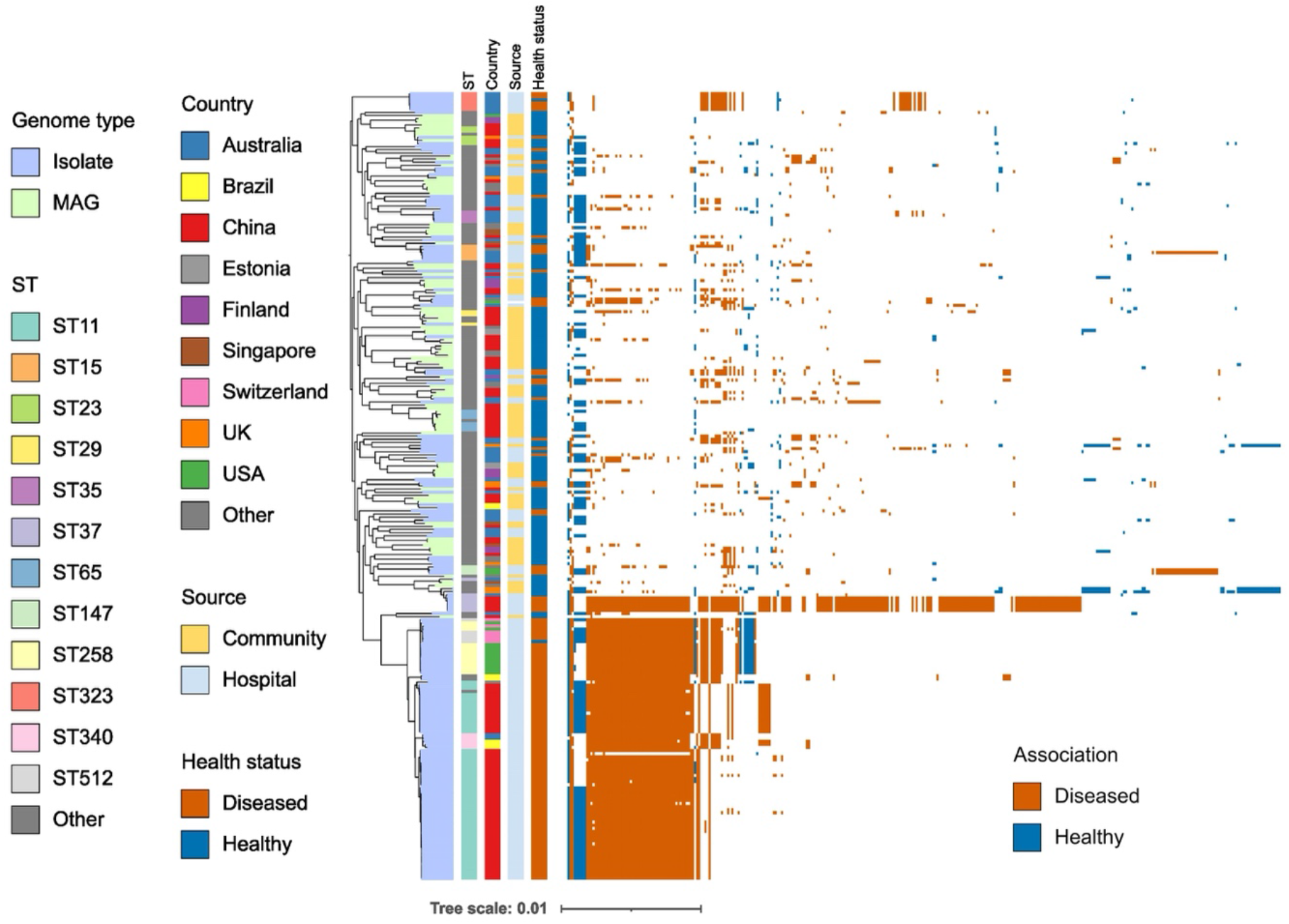
Phylogenetic tree and candidate genes linked to health or disease. Core-genome phylogenetic tree of *K. pneumoniae* genomes from carriage and disease (infection). The first four annotation blocks denote various metadata properties of the genomes, while the remaining layers to the right depict the distribution of all the significant genes associated with either carriage (healthy) or disease.

**Extended Data Figure 5.**
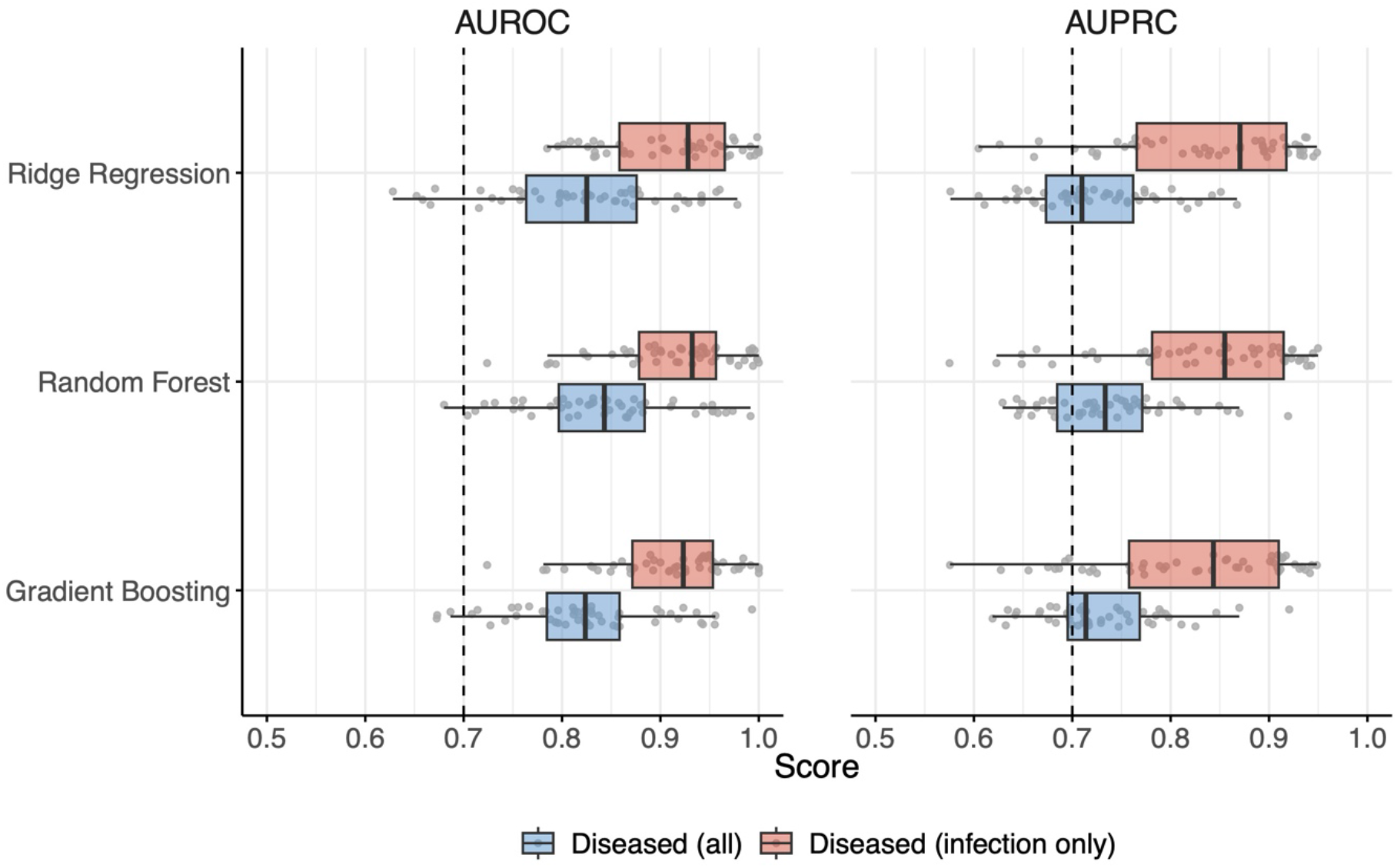
Classification of carriage and disease using supervised machine learning models. Performance of machine learning (ML) models distinguishing carriage- from disease-associated *K. pneumoniae*, considering either all disease genomes (blue) or just those from infection (red). Three supervised ML models were tested: ridge regression, random forest and gradient boosting. Each box in the plot represents the interquartile range (IQR) of the Area Under the Receiver Operating Curve (AUROC, left) or the Area Under the Precision-Recall Curve (AUPRC, right). The centre line within the box represents the median score. Whiskers are shown extending to the furthest point within 1.5 times the IQR from the box.

